# A genome-wide RNAi screen identifies MASK as a positive regulator of cytokine receptor stability

**DOI:** 10.1101/187286

**Authors:** Katherine H Fisher, Maria Fragiadaki, Dhamayanthi Pugazhendhi, Nina Bausek, Stephen Brown, Martin P Zeidler

## Abstract

In order for cells to sense and thus respond to their environment, they require transmembrane receptors, which bind extracellular ligands and then transduce this signal within the cell. A subset of receptors, with single-pass transmembrane domains are known as cytokine receptors and act via the Janus Kinase and Signal Transducer and Activator of Transcription (JAK/STAT) pathway. These receptors are essential for processes such as haematopoiesis, immune responses and tissue homeostasis. In order to transduce ligand activation, cytokine receptors must dimerise. However, mechanisms regulating their dimerisation are largely unknown. In order to better understand the processes regulating cytokine receptor levels, activity and dimerisation, we used the highly conserved JAK/STAT pathway in *Drosophila*, which acts via a single receptor, known as Domeless. We have performed a genome-wide RNAi screen in *Drosophila* cells, identifying MASK as a positive regulator of Domeless dimerisation and protein levels. We show that MASK is able to regulate JAK/STAT signalling both *in vitro* and *in vivo*. We go on to show that MASK is able to bind to Domeless via its Ankyrin repeat domains and alters the stability of the receptor. Finally, we extend our observations to the human homologue, ANKHD1, and demonstrate functional conservation, with ANKHD1 able to regulate JAK/STAT signalling and the levels of a subset of pathway receptors in human cells. Taken together, we have identified MASK as a conserved regulator of cytokine receptor levels, which may have implications for human health.

## Introduction

The ability to bind to extracellular ligands and transduce the resulting interaction across the plasma membrane represents the central biological function of cytokine receptors. Such receptors include the single-pass transmembrane proteins that ultimately stimulate the Janus Kinase and Signal Transducer and Activator of Transcription (JAK/STAT) pathway (Arbouzova and Zeidler, 2006, Vainchenker and Constantinescu, 2013). This group of receptors are typically homo‐ or hetero-dimerised with an extracellular structure consisting of multiple Fibronectin Type III repeats in which two of the distal repeats form a cytokine binding motif (Tanaka et al., 2014) (CBM; Fig 1A). On the C-terminal intracellular side, long, β-type cytokine receptors, such as GP130, Oncostatin M Receptor B (OSMRB), Thromobopoietin Receptor (TPOR) and the *Drosophila* receptor Domeless (Dome), contain juxta-membrane domains via which cytosolic JAK tyrosine kinases associate. By contrast, shorter, α-type receptors such as IL-6Rα, participate in the formation of ligand binding complexes but lack the intracellular domains bound by downstream pathway components (Heinrich et al., 2003).

**Figure 1.**
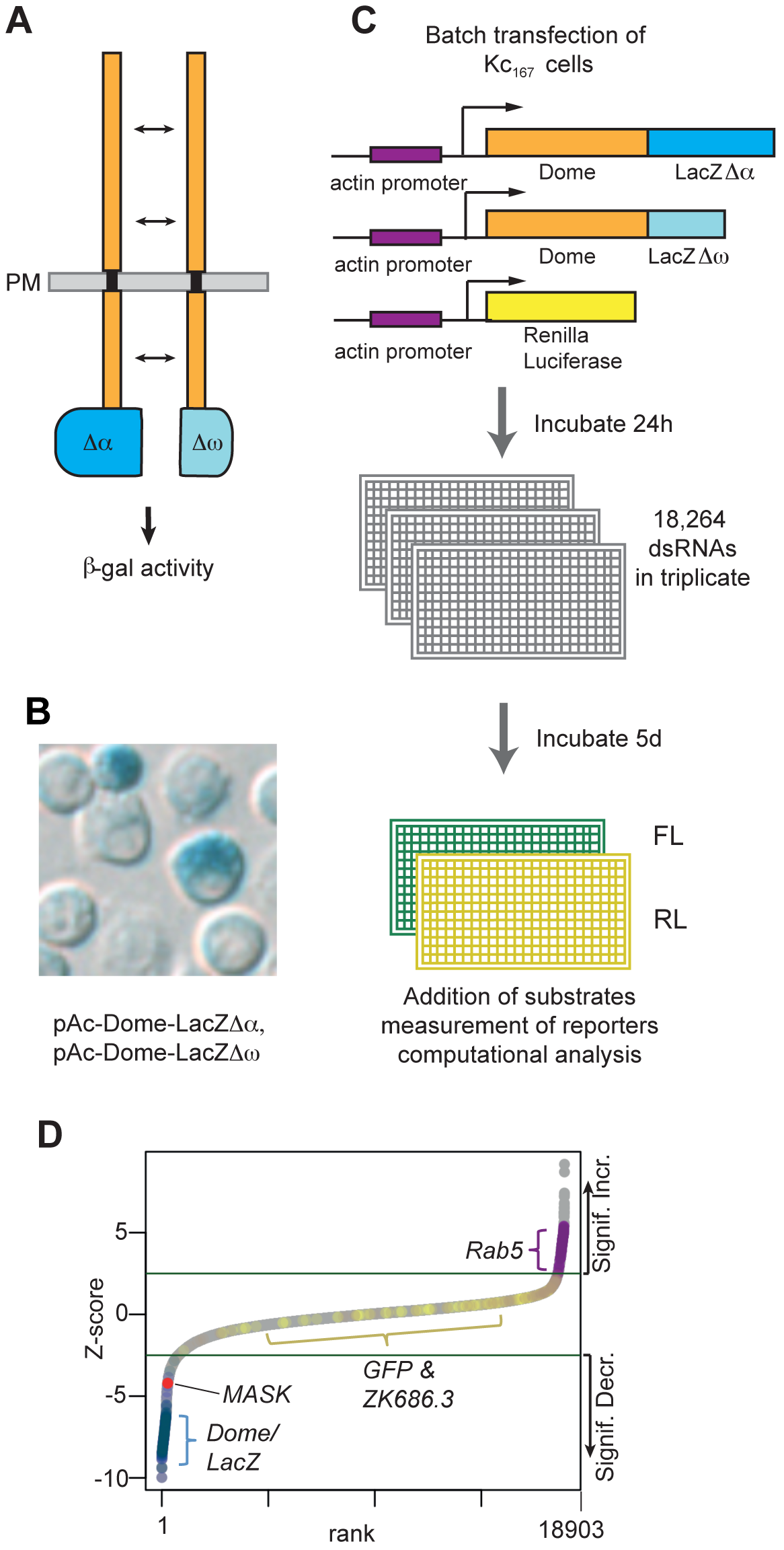
A split (3-galactosidase genome-wide RNAi screen for modulators of Dome dimerisation and levels. **A)** Schematic representation of the Dome-βgalΔα and Dome-βgalΔω complementation assay. PM = plasma membrane. **B)** *Drosophila* Kc_167_ cells transiently transfected with plasmids expressing the proteins shown in (A) show β-galactosidase activity by X-gal staining. **C)** Workflow of the genome-wide RNAi screen for modulators of Dome dimerisation and levels as undertaken in *Drosophila* Kc_167_ cells. **D)** Ranked Z-scores from the genome-wide RNAi screen. Green lines illustrate Z-score cut-offs of significant increase or significant decrease. Controls are shown with *MASK* highlighted in red.

Cytokine binding to the extracellular domains of a receptor complex then induces a conformational change, which either reorients a preformed dimer (Brown et al., 2005, Remy et al., 1999) or induces receptor dimerisation/oligomerisation (Thomas et al., 2011). In canonical JAK/STAT pathway signalling, this activation results in JAK auto-phosphorylation, followed by trans-phosphorylation of intracellular tyrosine residues in the receptor tail and recruitment of latent STAT molecules (Vainchenker and Constantinescu, 2013). These STATs are then themselves tyrosine phosphorylated, dimerise and translocate to the nucleus where they bind to palindromic DNA sequences in the promoters of pathway target genes and thus regulate gene expression.

In humans, JAK/STAT pathway signalling is mediated by 4 JAKs and 6 STATs, playing key roles both during embryonic development, adult homeostasis and multiple diseases (Arbouzova and Zeidler, 2006, Vainchenker and Constantinescu, 2013). These core components are themselves downstream of multiple receptors and ligands with cell-specific differences, redundancy and cross talk between pathway components making the dissection of signalling processes particularly challenging. For example, signalling by the pro-inflammatory cytokine IL-6, occurs via receptor heterodimers made up of the long β-type GP130 receptor and the shorter **α**-type IL-6R, with both membrane-bound and soluble forms of IL-6R able to form signalling competent complexes with GP130 to stimulate the downstream pathway (Tanaka et al., 2014). By contrast, the production of erythrocytes and platelets is dependent on homo-dimerised Erythropoietin Receptor (EPOR) and TPOR respectively, receptors which function upstream of JAK2 and STAT5 (Seubert et al., 2003). In addition, the maintenance of stem cell fate and the transduction of pro-proliferative signals involved in growth control and tissue remodelling are also frequently dependent on JAK/STAT pathway regulation. Examples include the requirement for the pathway in haematopoietic stem cell maintenance mediated by TPOR (Qian et al., 2007, Staerk and Constantinescu, 2012) and tissue remodelling within the mammary gland during and after lactation (Hughes and Watson, 2012).

Although the core JAK and STAT pathway components have been extensively studied, the regulatory processes controlling upstream pathway receptors are less well understood. One key mechanism regulating receptor levels at the plasma membrane is endocytosis. Originally considered as a mechanism to attenuate pathway signalling following activation (Liu and Shapiro, 2003), it is now clear that the endocytosis and trafficking of ligand: receptor complexes into endosomes, and continued pathway signalling from this internalised compartment, not only occurs, but is also frequently qualitatively changed (reviewed in (Cendrowski et al., 2016)). Although uncertainty remains, changes in the micro-environment within a maturing endosome such as reduced pH, trapping of the ligand, alterations / breakup of the receptor complex and changes to ligand:receptor affinities are all likely to occur (Kurgonaite et al., 2015). Ultimately, receptor recycling to the plasma membrane or destruction of the complex within the lysosome also change the levels of functional receptors (Chmiest et al., 2016, Gesbert et al., 2004).

In addition to the regulation of receptor protein levels, the formation of receptor complexes is also required for ligand binding and the formation of a functional signalling complex (Heinrich et al., 2003). However, it is clear that various mechanisms are associated with the dimerisation of distinct receptor populations. In the case of EPO, ligand binding has been shown to be sufficient to bring about receptor dimerisation (Boger and Goldberg, 2001) while the related receptor in *Drosophila,* Domeless (Dome), is dimerised *in vivo* via a ligand and JAK/STAT pathway-independent mechanism (Brown et al., 2003).

By contrast to the multiple components that make up the mammalian JAK/STAT pathway, the conserved pathway in *Drosophila melanogaster* is significantly less complex and features a single JAK and a single STAT-like molecule together with a single full-length receptor, Dome, and a single short antagonistic receptor, termed Latran [also known as Eye Transformer] (reviewed in (Zeidler and Bausek, 2013)). As is the case for vertebrate pathways, the *Drosophila* pathway is also involved in developmental processes, immunity, cellular proliferation and haematopoiesis (Arbouzova and Zeidler, 2006). In particular, conservation of pathway function is demonstrated by gain-of-function mutations in JAK kinases, which are associated with blood cell overproliferation in both fly models and human myeloproliferative disease (Harrison et al., 1995, Luo et al., 1997, Pasquier et al., 2014).

In order to better understand the processes regulating cytokine receptor levels, activity and dimerisation, we set out to exploit the low complexity of the *Drosophila* system by undertaking RNA-interference (RNAi)-based screens for regulators of the *Drosophila* receptor, Dome. A previous report indicated that JAK/STAT pathway activation downstream of the Dome receptor requires homo-dimerisation of the receptor (Brown et al., 2003). Furthermore, it showed that dimerisation is mediated by an as-yet unidentified ligand‐ and signalling-independent mechanism *in vivo.* In this study, we employed a molecular complementation assay utilising two truncated forms of the β-galactosidase enzyme termed Δα and a Δω (Rossi et al., 1997) and fused these to the cytosolic, C-terminal ends of the Dome receptor. Although enzymatically inactive in isolation, dimerisation of two Dome molecules brings together both a Δα and a Δω truncation, allowing molecular complementation and the reconstitution of β-galactosidase activity (Fig. 1A).

Here we present our use of such a molecular complementation assay to undertake RNAi screens for factors able to modulate Dome levels and/or dimerisation. We present our genome-wide analysis of this screen and go on to follow up by analysing the conserved, Multiple Ankyrin repeats and KH-domain containing protein, MASK. Using both biochemical and genetic approaches, we show that MASK promotes Dome dimerisation and stability and demonstrate that JAK/STAT pathway activity is reduced following MASK knockdown. We go on to demonstrate that MASK binds directly to the Dome receptor via its medial A2 cluster of Ankyrin repeats and stabilise the resulting complex. We show that the conserved human homologue, ANKHD1, is also able to regulate both JAK/STAT pathway activity and the stability of a subset of human cytokine receptors. This study therefore identifies a novel regulator of cytokine receptor levels providing insights into the regulation of this disease-relevant signalling pathway.

## Results

### A split β-galactosidase assay for monitoring receptor dimerisation

Genome-scale RNAi screening has previously identified multiple regulators of JAK/STAT transcriptional activity (Kallio et al., 2010, Müller et al., 2005). However, changes in gene expression do not provide insights into the molecular mechanisms via which regulators of the pathway act. We therefore modified an assay for Dome dimerisation using a split β-galactosidase (β-gal) complementation system (Brown et al., 2003), in which the coding region for the β-gal enzyme containing one of two inactivating deletions (termed Δα and Δω) was attached to the intracellular terminus of the Dome receptor (Fig. 1A). As previously demonstrated *in vivo* (Brown et al., 2003), each individual β-gal mutant is inactive in isolation and shows enzymatic activity only when co-expressed in the same cells (Fig. S1A). We adapted this technique for use in cultured *Drosophila* cells (Fig. 1B) and optimised a luminescent readout for β-gal enzymatic activity (Fig. S1A and Materials and Methods). Although designed to detect receptor dimers, our assay was also inherently sensitive to the absolute level of these dimers, since any changes in the amount of protein would also result in changes in β-gal activity.

### A genome-wide RNAi screen identifies modulators of Dome dimerisation and stability

We performed a genome-wide RNAi screen using a second generation, *in silico* optimised, double-stranded RNA (dsRNA) library targeting 97.9% of the *Drosophila* genome (Fig. 1C, Fig. S1B) and analysed the resulting >110k data points using best practice analysis techniques (Fisher et al., 2012). As expected, negative controls (targeting *GFP* or the *C.elegans* gene *zk686.3)* did not significantly affect our assay while knockdown of the endocytic trafficking component *rab5* increased levels of dimerised Dome, consistent with previous reports (Stec et al., 2013, Vidal et al., 2010). Conversely, knockdown of either *dome* itself or *lacZ* strongly decreased β-gal activity (Fig. 1D; Fig. S1B). Using techniques previously developed for similar genome-scale screens (Fisher et al., 2012, Müller et al., 2005), we first identified hits from three replicates of initial screening and subsequently retested these in secondary re-screens (Table S1). Based on both primary and secondary screening, 43 candidates with consistent and robust Z-scores were selected for further analysis (Table 1; see Materials & Methods for precise selection criteria). Previous work undertaken *in vivo* suggested that ligand expression is not sufficient for Dome dimerisation (Brown et al., 2003). To test this finding in the context of our 43 candidates, we repeated the original Dome dimerisation assay in the presence of co-expressed pathway ligand and found that most (79%, 34/43) of the original hits reproduced their effects (Table 1). In addition, it has also been shown that Dome can form hetero-dimers with the short negatively acting pathway receptor Latran (Lat), and that Lat can also form homodimers with itself (Fisher et al., 2016, Makki et al., 2010). We therefore set up cell based assays to test for Dome:Lat heterodimer and Lat:Lat homodimer formation and used this to test the 43 candidate genes. We found that 90% (36/40) of candidates affect Dome:Lat and Lat:Lat dimers, with 31 of these common to both (Table 1).

**Table 1.**
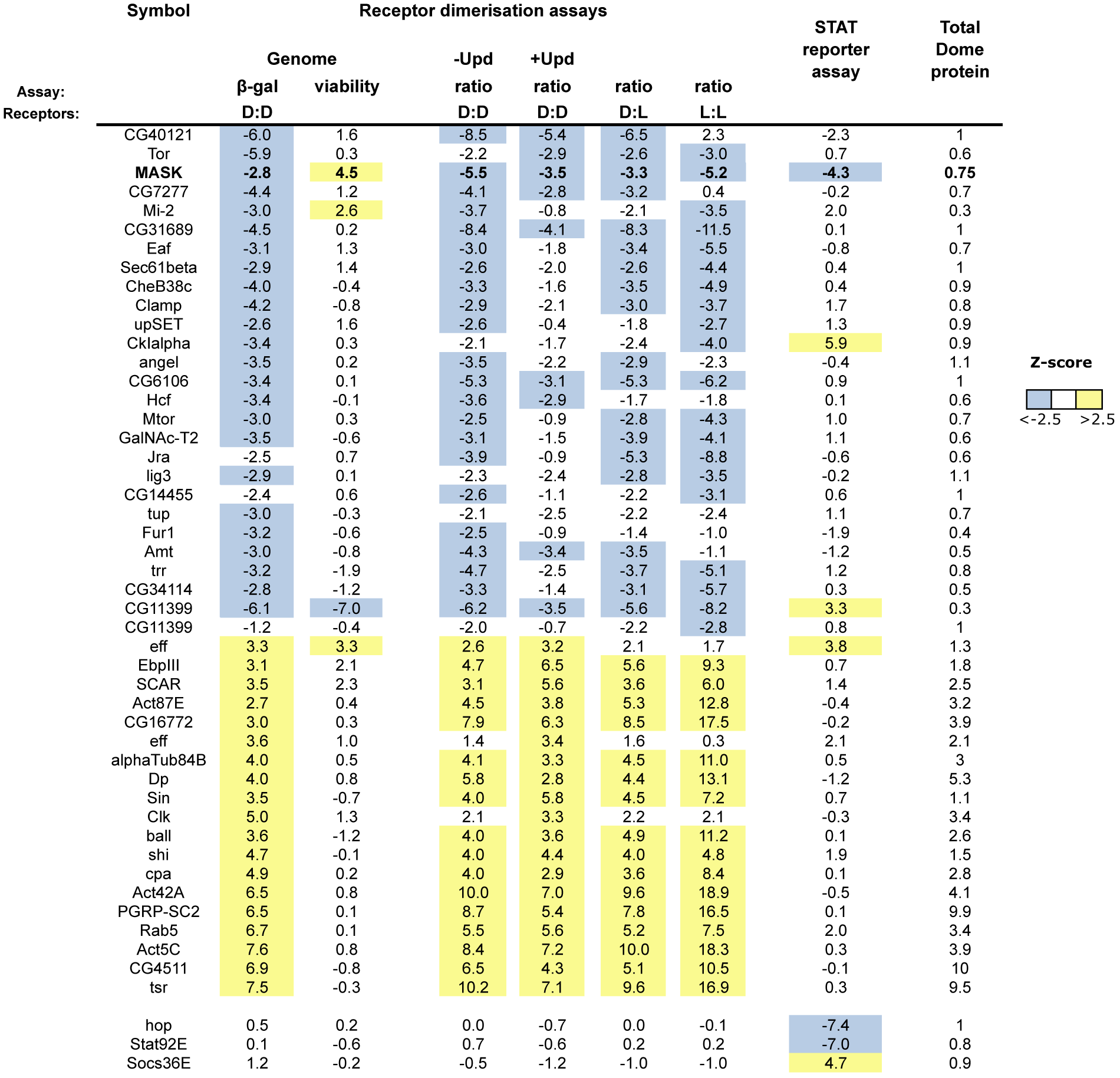
Secondary screens confirm 43 robust hits. After secondary screening with Dome receptor assay, 43 candidates were found to be reproducible hits (see Table S1 and Materials and Methods for further details). Two genes are listed twice, since independent dsRNAs for these were present within the genome plates and taken through to secondary screens. Rab5 was also identified, even though this was used as a positive control. To prove the effectiveness of the STAT reporter assay, we have included results from knockdown of hop, STAT92E and SOCS36E. Results for MASK are shown in bold. Numbers for receptor dimerisation and STAT reporter assays are shown as Z-scores. Dome protein levels are shown as fold-change relative to controls.

Although our molecular complementation assay requires receptor dimerisation to produce β-gal enzymatic activity, changes in signal can also be a consequence of changes in overall receptor levels. In order to distinguish between those hits that promote or inhibit dimerisation and those that simply change protein levels, we next sought to quantify total protein levels using an independent technical approach. We therefore used quantified Western blotting undertaken in triplicate (see Supporting Information for assay design) to examine the effects of the 43 genes on overall Dome protein levels. Of the candidates tested, 31 altered Dome protein levels by at least 25% (Table 1, also see Fig. S1C-D for examples) while the remaining 13 appear to change dimerisation without affecting overall protein levels.

Given the changes in Dome dimerisation and protein levels we also assessed the effect of our hits on JAK/STAT dependent transcription using the *6x2xDrafLuc* reporter (Müller et al., 2005). Surprisingly, while knockdown of some genes clearly affected JAK/STAT transcriptional activity, a large proportion of the dsRNAs had little or no effect on the *6x2xDrafLuc* reporter (Table 1). This rather unexpected result suggests that either the levels of dimerised receptor are not rate limiting in this system, or that alternative regulatory pathways are able to compensate for changes in Dome dimer activity.

We next undertook an analysis of our 43 candidates to identify gene ontological terms disproportionately enriched or depleted relative to the whole *Drosophila* genome (Mi et al., 2017). This identified strong overrepresentation of genes involved in endocytosis (GO:0006897), actin cytoskeleton (GO:0015629), and cellular component morphogenesis (GO:0032989).

One striking GO group identified were actin-related proteins initially identified as strong hits, a group of hits which also resulted in significant up-regulation of Dome protein levels (Table 1 and Fig. S1E). Upon further examination by qPCR, we found that RNAi-mediated knockdown of *Act42A* in our cells resulted in a significant increase in the transcription of Dome construct transfected into our cells and expressed by an *actin5c*-derived promoter (Fig. S1F). Given that this result indicated the existence of a feedback loop regulating the Actin promoter used in this construct, Actin-related genes were classified as non-specific in this assay.

### MASK regulates levels of dimerised Dome

Throughout the multiple rounds of screening and secondary assays undertaken, RNAi targeting MASK consistently generated strong effects on the dimerisation assay, receptor levels and JAK/STAT pathway transcriptional activity (bolded in Table 1). We therefore, set out to better investigate the mechanisms underlying this activity.

In order to confirm the screen-based identification of MASK, we retested its effect using an alternative dimerisation assay. Using co-immunoprecipitation of differentially epitope tagged Dome molecules followed by quantification of western blots, we found that knockdown of MASK was sufficient to reduce levels of dimerised Dome by 50% (± 10%, p<0.013, n=3) (Fig. 2A). However, this change in dimerisation was also accompanied by an approximate 25% decrease in the steady state level of Dome protein detected following MASK knockdown (Fig. 2B). As such, knockdown of MASK resulted in both the destabilisation of Dome dimers and also a reduction in the absolute levels of Dome itself.

**Figure 2.**
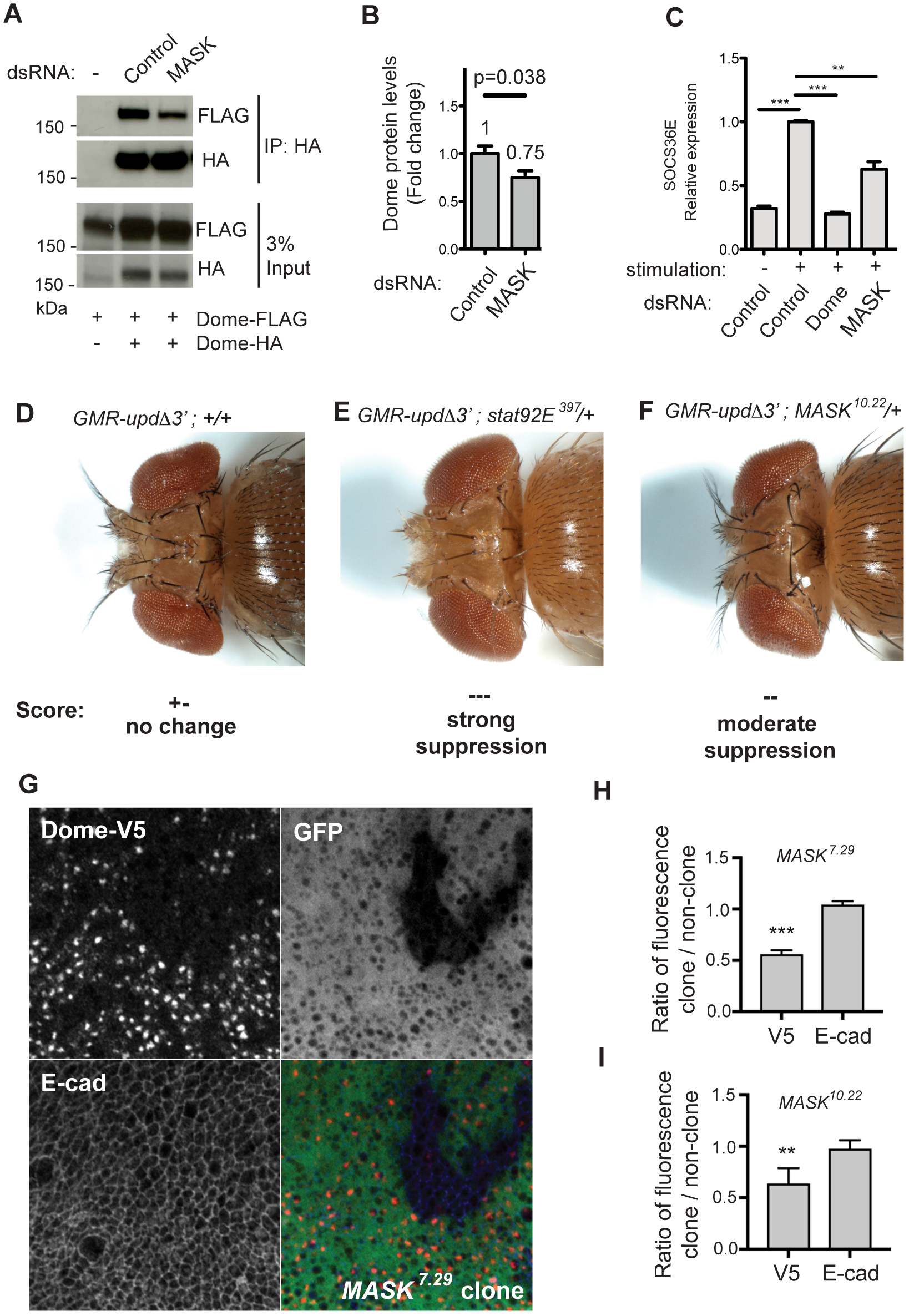
MASK regulates pathway activity and receptor levels *in vivo*. **A)** Co-expression of Dome-HA and Dome-FLAG followed by Dome-HA immunoprecipitation in *Drosophila* Kc_167_ cells. Levels of co-precipitated Dome-FLAG are modulated by treatment with the MASK dsRNA. **B)** Quantification of steady state Dome-FLAG protein levels expressed by Kc_167_ cells after knockdown of MASK. Number indicates fold change, error bars show standard deviation, p-values are indicated (n=3). **C)** Expression of the JAK/STAT pathway target gene *SOCS36E* following Upd2 ligand stimulation and treatment with indicated dsRNAs. Error bars show standard deviation (n=3), ** *p* < 0.01 *, *** p <* 0.001. **D-F**) Dorsal view of eye overgrowth phenotypes caused by ectopic Upd ligand expression driven by *GMR-UpdΔ3’*. Loss of one copy of *STAT92E* or *MASK* suppresses overgrowth. **G)** Mitotic clones of *MASK*^*7.29*^ caused a reduction in *tubulin-GAL4* driven UAS‐Dome-V5 (red) fluorescence, whereas E-cadherin (blue) levels were unaffected. Clones were identified using loss of native GFP (green). **H-I**) Quantification of Dome-V5 and E-cad levels in *MASK*^*729*^ (H) or *MASK*^*10.22*^ (I) mutant clones. Ratios of fluorescence intensity inside clones and in nearby twin-spots were taken to control for variations across discs. Measurements were averaged over **>**4 discs with at least 2 clones per disc. ** *p <* 0.01, *** *p* < 0.001 (One-sample t-test with expected mean of 1).

Consistent with the decrease in levels and dimerisation of receptor, we also found that Upd-induced JAK/STAT transcriptional activation was reduced following knockdown of MASK, both at the level of a luciferase-based JAK/STAT-sensitive reporter (Table1) and also via the reduction in transcription of the STAT92E target gene, *SOCS36E* (Fig. 2C). This result was confirmed using two independent dsRNAs, each of which reduced MASK mRNA by >70% (Fig. S2A), and strongly reduced pathway-induced transcription (Fig. S2B). The requirement for MASK in JAK/STAT signalling was further demonstrated using two additional independent STAT92E reporter assays, each of which was strongly and significantly reduced following MASK knockdown (Fig. S2C). Taken together, these findings confirm that MASK functions as a positive regulator of the *Drosophila* JAK/STAT pathway.

Previous reports have identified MASK as a regulator of the Ras/Raf and Hippo/Warts pathways (Sansores-Garcia et al., 2013, Sidor et al., 2013, Smith et al., 2002), we examined the effects of silencing known components of the Ras (*csw, ras85D, ras64B, raf*) and Hippo (*hpo, wts, yki*) pathways in order to identify potential pathway cross-talk with our JAK/STAT pathway assays. Analysed as Z-scores relative to the original genome-wide screen dataset, neither Dome dimerisation/stability (Fig. S2D) nor STAT92E transcriptional activity (Fig. S2E) were significantly affected by knockdown of any of the Ras or Hippo pathway components tested. This suggests that MASK is acting directly on the JAK/STAT pathway.

### MASK regulates JAK/STAT signalling *in vivo*

In order to support the cell and RNAi-based data, we also undertook *in vivo* JAK/STAT pathway assays using previously characterised loss-of-function MASK alleles (Smith et al., 2002). Ectopic JAK/STAT pathway activation is sufficient to drive over-proliferation within the developing eye imaginal disc, a process that is sensitive to downstream JAK/STAT pathway activity (Fig. 2D-E; (Bach et al., 2003, Mukherjee et al., 2006)). Using this test, we found that JAK/STAT pathway induced eye overgrowth was markedly reduced in genetic backgrounds heterozygous for independent MASK loss-of-function alleles (Fig. 2F and Fig. S2F-H).

We next explored whether MASK was required to maintain Dome protein levels *in vivo*. Since existing MASK mutant alleles are embryonic lethal, we induced mitotic clones of either the hypomorphic allele *MASK^7^*^.29^ or the null *MASK^10.22^* allele in developing wing discs (Fig. 2G-H). In the absence of reliable antibodies against Dome, we ubiquitously expressed epitope-tagged Dome-V5 throughout the wing disc using *tubulin-GAL4* (Fig. 2G). As observed previously (Makki et al, 2010), Dome was found to accumulate in intracellular vesicles, with weaker staining also observed at the plasma membrane (not shown). Although *MASK* mutant clones proliferate poorly and are therefore relatively small, levels of Dome detected in mutant areas are significantly lower than in surrounding wild type tissue (Fig. 2G-I). When Dome levels inside clones (which are identified by their lack of GFP marker expression) were quantified relative to equivalent neighbouring, wild-type regions, a significant reduction in Dome-V5 levels was observed for both *MASK* alleles (Fig. 2H,I). By contrast the single pass transmembrane protein E-cadherin (E-cad), which is not a JAK/STAT pathway receptor, is unaffected by loss of MASK. Given that transcription of Dome in this experiment is driven via a uniformly expressed heterologous *tubulin* promoter, it is likely that changes in Dome levels are a function of reduced protein rather than a change in gene expression. These results suggest that MASK acts as a positive regulator of the JAK/STAT pathway *in vivo* in *Drosophila* and is able to modulate pathway receptor levels.

### MASK can physically associate with Dome

Given the ability of ankyrin repeats to mediate protein:protein interactions (Bennett and Chen, 2001, Michaely et al., 1999) and given the ability of *Drosophila* MASK to modulate Dome receptor levels, we reasoned that MASK proteins may directly bind to cytokine receptors. We therefore utilised constructs encoding each of the ankyrin repeat domains and the KH domain of MASK (Sansores-Garcia et al., 2013), expressed these in *Drosophila* Kc_167_ cells (Fig. 3A) and tested these for binding to Dome. We found that the MASK-A2 and, more weakly, the MASK-A1 ankyrin repeat domains were able to co-precipitate Dome-FLAG while no interaction was detected with the MASK-KH region (Fig. 3B). This interaction was found to be reciprocal, with a FLAG-tagged construct containing both ankyrin repeat domains (MASK-A1A2) being immunoprecipitated with Dome-HA (Fig. 3C, schematic of this construct in 3A).

**Figure 3.**
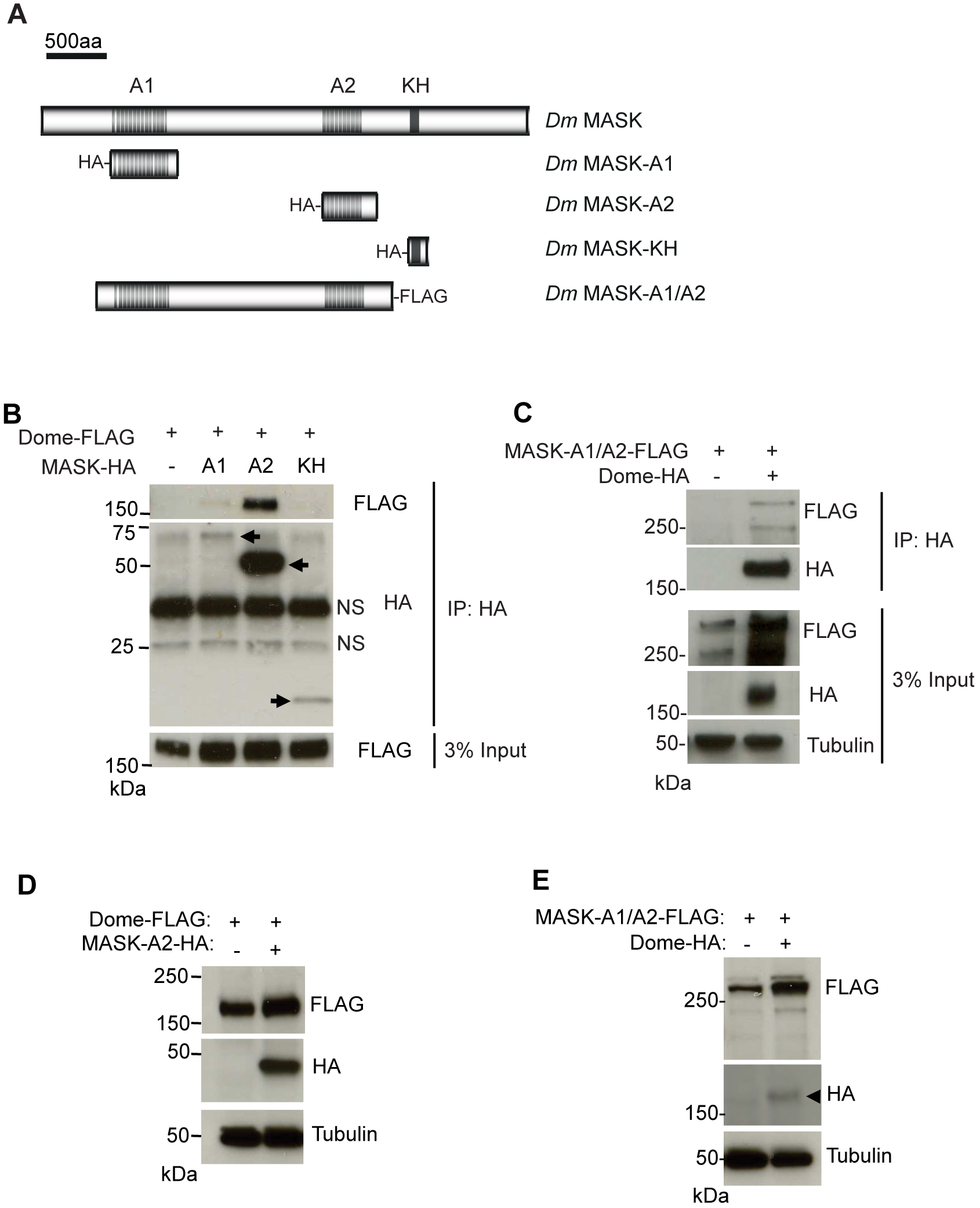
MASK physically associates with Dome. **A)** Schematic representation of *Drosophila* MASK protein and constructs used in this study. **B)** Immunoprecipitation of the indicated HA-MASK constructs from Kc_167_ cells also expressing Dome-FLAG. Dome-FLAG is co-immunoprecipitated with HA-MASK-A1 and HA-MASK-A2. Levels of Dome-FLAG present in the input lysate are shown. NS = non-specific band. **C)** Co-precipitation of MASK-A1/A2-FLAG following immunoprecipitation of Dome-HA. Levels of MASK-A1/A2-FLAG, Dome-HA and α-Tubulin present in the total Kc_167_ cell lysates are shown. **D)** Steady state levels of Dome-FLAG expressed in *Drosophila* Kc_167_ cells are increased following the co-expression of HA-MASK-A2. Levels of α-Tubulin indicate loading parity. **E**) Steady state levels of MASK-A1/A2-FLAG expressed in *Drosophila* Kc_167_ cells are increased following the co-expression of Dome-HA. Levels of α-Tubulin indicate loading parity.

These results suggest that MASK is able to form a physical complex with Dome and suggest that this interaction occurs via its ankyrin repeat domains.

### Increased MASK levels leads to raised receptor levels

Given the negative effect of RNAi-mediated MASK knockdown on receptor stability, we tested whether increased levels of MASK fragments might have the opposite effect. In contrast to loss-of-function experiments, we found that expression of the second MASK ankyrin repeat cluster (MASK-A2; Fig. 3A; (Sansores-Garcia etal., 2013)) was sufficient to increase the steady state levels of Dome in Kc_167_ cells (Fig. 3D). Consistent with the reduction in Dome levels detected following MASK knockdown (Table 1; Fig. 2B), over expression of exogenous Dome was also sufficient to reciprocally stabilise MASK-A1A2, a construct comprising both the first and second ankyrin repeat clusters (Fig. 3E). These results support the initial finding that MASK is a positive regulator of Dome stability and when taken together with the physical interactions between Dome and MASK, suggest that Dome:MASK association forms a stabilised protein complex.

### Conservation of MASK function in human cells

Since MASK has been evolutionarily conserved between humans and *Drosophila* at the primary sequence level (Fig. 4A), we tested whether its function in modulating JAK/STAT signalling is also conserved. To test the effects of knocking down the closely related ANKHD1 on the human JAK/STAT pathway, we used qPCR in HeLa cells to detect the ligand-dependent increase in the mRNA levels of the pathway target gene, *SOCS3* (Murray, 2007). As expected, mRNA levels of *SOCS3* were strongly decreased following silencing of JAK2 and STAT3 while siRNA-mediated silencing of ANKHD1 (whose efficiency is shown in Fig. S3A) was also sufficient to significantly reduce expression (Fig. 4B). Consistent with this, ANKHD1 knockdown was also sufficient to significantly reduce the OSM-stimulated phospho-STAT1 and phospho-STAT3 levels in this system (Fig. 4C and Fig. S3B).

**Figure 4.**
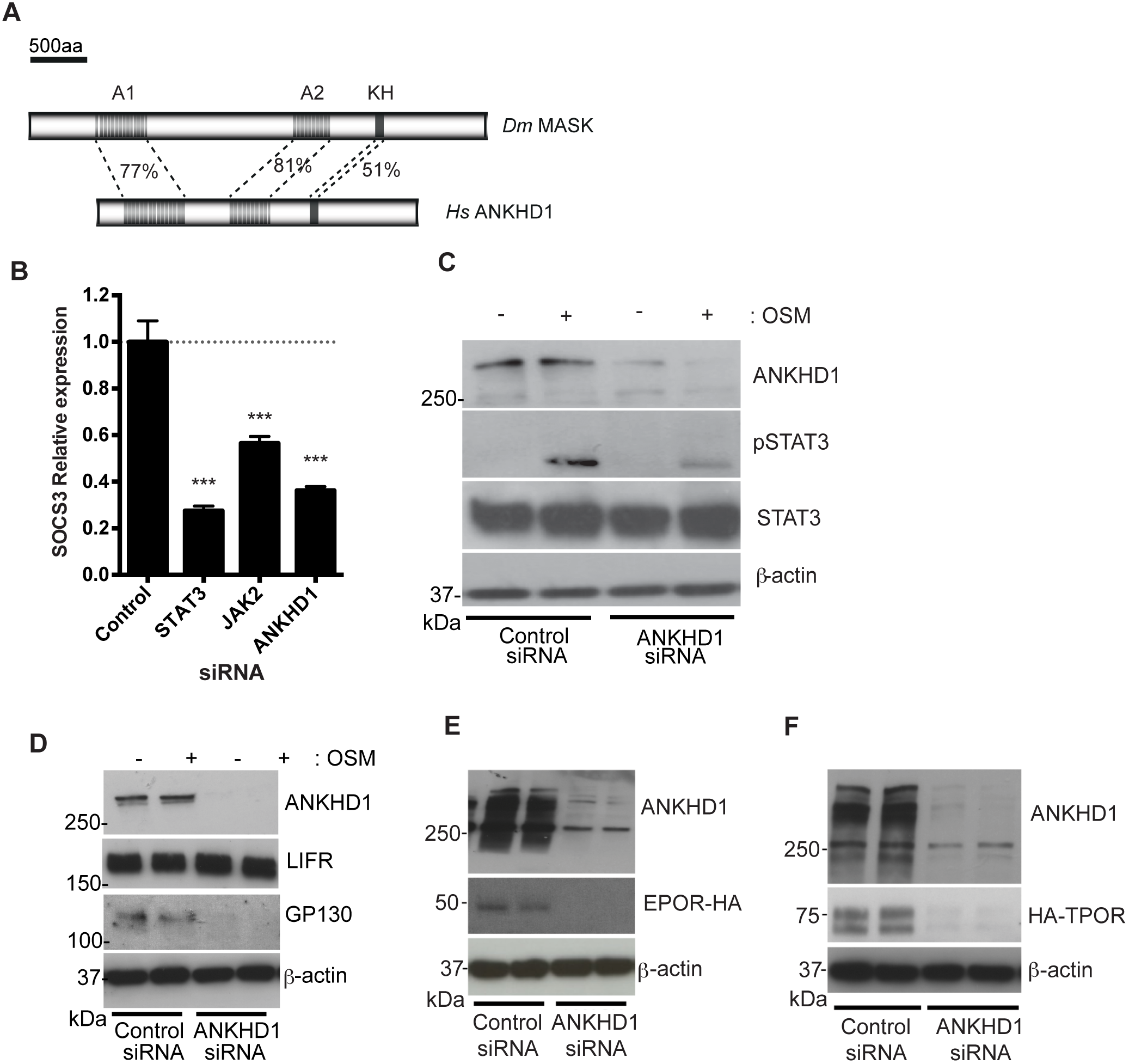
MASK function is conserved to human cells. **A)** Schematic representation of protein structure of *Drosophila* MASK and human ANKHD1 with % identity between sequences. **B)**HeLa cell extracts treated with siRNA targeting ANKHD1 have reduced phospho-STAT3 upon OSM stimulation, which total STAT3 levels are unaffected. Blots confirm knockdown of ANKHD1 levels. **C)** *SOCS3* mRNA expression in HeLa cells transfected with indicated siRNA and following OSM stimulation as indicated. **D)** A representative blot of HeLa cells treated with control siRNA or siRNA targeting ANKHD1 in (n=3). Silencing of ANKHD1 leads to a loss of both ANKHD1 protein and endogenous GP130 protein. By contrast, levels of LIFR are not changed. β-actin levels are unaffected. **E-F**) HeLa cell extracts treated with siRNA targeting ANKHD1 have reduced HA-EpoR (G) and TpoR-HA (H) compared to controls. Blots confirm knockdown of ANKHD1 levels.

Given that knockdown of *Drosophila* MASK led to a reduction in Dome protein levels (Fig. 2B, 2G) we tested cytokine receptor levels in human cells. Strikingly, while knockdown of ANKHD1 had no detectable effect on Leukaemia Inducible Factor receptor (LIFR), it led to the almost complete loss of the endogenous glycoprotein 130 (GP130), the long cytokine receptor central to IL6-class cytokine receptor complexes (Fig. 4D; (Heinrich et al., 2003)). This change in GP130 protein level may be the consequence of either changes in protein stability, mRNA stability or transcriptional regulation. In order to provide insight into the basis of receptor stabilisation we tested the ability of ANKHD1 to alter the levels of HA-tagged TPOR and EPOR expressed from a CMV promoter in HeLa cells in the presence or absence of ANKHD1. Both receptors were greatly reduced following treatment with ANKHD1 siRNA (Fig. 4E,F). Since these receptors were expressed from a constitutive promoter, they are unlikely to be affected by changes in transcriptional control, further supporting a model in which ANKHD1 functions at a post-transcriptional level.

Taken together, these results suggest that the human homologue of MASK, ANKHD1, also acts as a positive regulator of JAK/STAT signalling and modulates the levels of a subset of human cytokine receptors.

## Discussion

The development, and maintenance of multicellular life is absolutely dependent on the ability of cells to communicate with one-another - a process that requires transmembrane receptor molecules. In this report we have undertaken a screen to identify the factors involved in the dimerisation and stability of the *Drosophila* receptor associated with JAK/STAT pathway activation. This single pass, trans-membrane receptor, termed Dome, forms homo-dimers in a spatially and temporally restricted manner during embryonic development. This dimerisation is required for downstream signalling, but is unaffected by the presence of the pathway ligand Unpaired (Brown et al., 2003). More recently, a related, but shorter receptor, termed Latran, was identified which acts negatively to down-regulate JAK/STAT pathway signalling (Kallio et al., 2010, Makki et al., 2010). Strikingly, Latran has also been shown to be able to form both homo-dimers and hetero-dimers with Dome (Fisher et al., 2016, Makki et al., 2010) with the formation of signalling-incompetent Dome:Lat heterodimers thought to represent the mechanism of negative regulation (Fisher et al., 2016). However, while the receptors themselves have been characterised, the mechanisms mediating receptor dimerisation required to generate a signalling-competent complex are unknown. As such, the data reported here represents the first comprehensive description of the components required for this process.

One caveat of the screen design presented is the fact that the β-gal activity measured is influenced by both the efficiency of dimer formation and the stability/levels of Dome protein. In order to differentiate between these two influences, we undertook secondary screens using semi-quantitative Western blotting to assess protein levels. In this way we differentiated between those genes modulating dimerisation, those modulating protein levels and those that regulate both aspects. Based on this insight, the identification of hits that change β-gal activity and NOT protein levels (e.g. *sec61β* and *CG6106)* suggest that failure to dimerise does not inherently affect protein stability. By contrast, hits such as MASK that change both dimerisation and protein levels may be affecting both processes, although it is also possible that the loss of receptor stability following MASK knockdown may result in the breakdown of existing dimers as a prelude to protein destruction.

While transmembrane proteins destined for insertion into the plasma-membrane are processed via conserved ER and Golgi pathways, it is clear that knockdown of MASK does not globally affect the production and/or trafficking of all membrane spanning proteins. Rather, the requirement for MASK is specific to a subset of transmembrane proteins affecting Dome but not E-cadherin in *Drosophila* (Fig. 2G-I), and GP130, EPOR and TPOR, but not LIFR, in human HeLa cells (Fig. 4D-F). However, the basis of this specificity remains to be determined.

In order to obtain a mechanistic insight into the function of MASK, we also undertook a structure-function analysis of MASK itself. This showed that MASK and Dome form stable physical interactions as shown by reciprocal co-immunoprecipitation, with this interaction being primarily mediated by the second, central A2 group of ankyrin domains present in MASK (Fig. 3B). Furthermore, we also show that the overexpression of the MASK-A1/A2 region is able to stabilise Dome levels (Fig. 3E), suggesting that MASK:Dome complexes [and possibly MASK:Dome:Dome complexes] may be inherently more stable than Dome alone.

Finally, we went on to demonstrate that ANKHD1, the human homologue of *Drosophila* MASK, also shares conserved functions, including a role in the regulation of JAK/STAT pathway signalling, with the levels of at least some receptors being strongly affected by changes in ANKHD1 levels (Fig. 4). In humans, *ANKHD1* has a paralogue on chromosome 4, named *ANKRD17* (ankyrin repeat domain 17) (Sansores-Garcia et al., 2013, Sidor et al., 2013) with the two proteins sharing 71% identity, with greater sequence similarity in the regions of ankyrin repeats and the KH domain (Poulinetal.,2003). Strikingly, ANKRD17 has been demonstrated to physically interact with receptors involved in the innate immune response, and plays a role in the release of cytokines (Menning and Kufer, 2013) and interferons (Wang et al., 2012). These findings serve to support our own data and suggest that ANKHD1 and ANKRD17 may also be acting to regulate receptor stability and dimerisation in humans.

Taken together, we present the first systematic screen, which we are aware of, to identify the factors responsible for the dimerisation of a JAK/STAT pathway receptor. We characterise one of these hits, MASK, and show that it regulates JAK/STAT pathway activity and forms a complex with the pathway receptor. We show that MASK is required to maintain the stability of Dome both *in vivo* and in cells and may well also play a role in receptor dimerisation. Finally, we demonstrate the evolutionary conservation of the MASK homologue ANKHD1 at the sequence and functional levels. As such, this work provides a valuable insight into this aspect of JAK/STAT pathway and highlights a novel level of regulation of this important and disease-relevant pathway.

## Materials and Methods

### Cell culture and biochemistry

*Drosophila* Kc_167_ cells were maintained according to standard procedures (Fisher et al., 2012). HeLa cells were maintained in DMEM supplemented with 10% serum. Plasmid transfections were carried out using Effectene (Qiagen) according to manufacturer’s instructions. Reverse transfections with siRNA were carried out using Lipofectamine RNAiMAX (Invitrogen) using 10nM final concentrations of single siRNAs (Dharmacon), targeting ANKHD1 (D-014405-01 or D-014405-02) where comparable results in terms of knockdown efficiency and reduction in JAK/STAT pathway activity were seen for both, or non-targeting siRNA as a control, D-001210-01. Stimulation of mammalian JAK/STAT pathway was carried out using human recombinant oncostatin M, (295-OM-010, R&D systems) at a final concentration of 10ng/ml for 20 minutes. Immunoprecipitation experiments were carried out as previously described (Stec et al., 2013). Proteins were separated on 4-15% TGX SDS-PAGE precast gels (Bio-Rad) and transferred to nitrocellulose membranes.

*Drosophila* RNAi screen hits were assessed for their effects on Dome protein levels. Kc_167_ cells were batch transfected with Dome-FLAG, incubated for 24h, then split into 24-well plates with 4μg dsRNA. After 5d RNAi treatment, cells were lysed as described. Lysates were boiled in 2x Laemmeli sample buffer and analysed by western blotting. FLAG/tubulin fold-changes were calculated for each RNAi condition in comparison to the average of three negative controls per gel. Each screen hit was analysed blind in duplicate.

### Genome-wide RNAi screening

The genome-wide SRSFv1 library, in 384-well format, was used as previously described (Fisher et al., 2012). Controls were manually added into empty wells (250ng dsRNA in 5ul water): *GFP* and the *C.elegans* gene baring no sequence homology in *Drosophila*, *zk686.3,* were used as baseline controls; technical controls targeting transfected plasmids were *dome, LacZ* and *RLuc*, and *Rab5* was used as a positive control. Genome-wide screening was carried out in biologically independent triplicates. Kc_167_ cells were batch-transfected in T75 flasks with 4μg *pAc-Dome-LacZ-Δα, pAc-Dome-LacZ-Δω* and *pAc-RLuc* and incubated for 24h. Cells were pooled in serum-free media, and 15,000 cells seeded per 384-well. After 1h, media was added to a final 10% serum concentration. After 5d cells were assayed for β-galactosidase activity using β-glo Assay System (Promega), which involves a Firefly luciferase reactions (FL), followed by measurement of Renilla luciferase (RL) activity as a viability control. Luciferase activities were measured on a Varioskan plate reader (Thermo).

### Data analysis

Firefly and Renilla luciferase values for each well were processed using the CellHTS2 Bioconductor package (Boutros et al., 2006). Values were median centred to normalise for plate-to-plate variation. Ratios of luciferase (FL/RL) were used to calculate the robust Z-scores, which were considered significant ≥2.5 or ≤-2.5. Individual FL and RL values were also assessed, since they were not always linear with respect to one another. Secondary analyses were carried out with newly synthesised dsRNAs and hits were considered significant at the less stringent ≥+2 or ≤-2. Forty-three robust hits were selected at this stage and sequenced to confirm correct target genes.

### Drosophila genotypes

Figure 2

D) *w, GMR-updΔ3’* / *w*^*1118*^

E) *w, GMR-updΔ3’* / *+;; stat92E*^*397*^/*+*

F) *w, GMR-updΔ3’* / *+;; MASK*^*1022*^/*+*

G,H) *w UbxFLP;;UAS-Dome-V5 FRT82 MASK*^*729*^ / *tub-GAL4 FRT82 Ubq-GFP*

I) *w UbxFLP;;UAS-Dome-V5 FRT82 MASK*^*10.22*^ / *tub-GAL4 FRT82 Ubq-GFP*

MASK alleles were a gift of M Simon (Smith et al., 2002).

### *Drosophila* phenotypes

Eye overgrowth assays were double blind scored alongside *stat92E* and *w^1118^* out-crosses (n>20 flies per genotype with >2 repeats). Adult flies were photographed using a Nikon SMZ1500 stereo-microscope and Nikon Elements extended depth of focus software package.

Wing discs were dissected from wandering 3^rd^ instar larvae raised at 25 °C. Inverted carcasses were fixed in 4% paraformaldehyde in PBS for 20min, blocked and incubated in primary antibodies overnight at 4°C. Tissues were washed in PBS containing 0.2% Triton X-100 (PBST) and incubated in secondary antibodies overnight at 4°C. After washing, discs were mounted in mounting media and imaged on Nikon A1R GaAsP confocal microscope using a 60x NA1.4 apochromatic lens, with a pixel size of 70 nm, and the pinhole was set to 1.2 AU.

### Antibodies

For western blot all primary antibodies were used at 1:1000 dilutions: ANKHD1 (Sigma, SAB1302423 and HPA008718), β-actin (Abcam, ab8226), GP130 (Cell Signaling, 3732 & Abcam ab34324), pSTAT1 (Cell Signaling 9167), STAT3 (Cell Signaling, 12640), pSTAT3 (Cell Signaling, 9145), FLAG (mouse M2, Sigma), HA (rat 3F10, Roche), *Drosophila* α-tubulin (mouse DM1A, Sigma). For immunohistochemistry, concentrations were E-cadherin (dCAD2, rat, DSHB, 1:20), V5 (mouse, Invitrogen, 1:500).

### Cloning of expression constructs

Dome-LacZ -Δα and ‐Δω fragments were cut from pUAST vectors (Brown et al., 2003) and ligated into pAc5.1 vector (Invitrogen) using KpnI and XbaI restriction sites (partial digestion of KpnI sites used for Δω). pAc-Dome-FALG and pAc-Dome-HA were described in (Stec et al., 2013). *Drosophila* MASK-A1/A2 was PCR amplified from cDNA clone LD31446 (DGRC). Gateway cloning of PCR product was carried out using the pENTR/D-TOPO Cloning Kit (Invitrogen) and introduced into the pAWF vector *(Drosophila* Gateway Vector Collection) using Gateway LR Clonase II Enzyme Mix (Invitrogen), according to the manufacturer’s instructions. HA-MASK constructs were a gift from G/ Halder (Sansores-Garcia et al., 2013).

### Quantitative real-time PCR

Total RNA was extracted from cells using TRIZOL Reagent (Invitrogen) following manufacturer’s instructions. Synthesis of cDNA was carried out using High Capacity RNA-to-cDNA Kit (Applied Biosystems) from 2 μg total RNA. To confirm gene knockdown by RNAi or to measure levels of target gene expression, qPCR was carried out using SsoAdvanced SYBRGreen Supermix (BioRad) on a CFX-96 Touch new generation Real-Time PCR Detection System (BioRad). Change in expression levels between experimental conditions was calculated compared to housekeeping gene expression (either *Drosophila* RpL32 or human β-actin) using the ΔΔC_T_ method (Bina et al., 2010). Statistical analysis was carried out using one-way ANOVA tests in Prism (Graphpad). Primers are listed in Table S2. TAQMAN qPCR probes were designed for multiplexing (IDT oligo).

## Acknowledgements

The authors would like to thank Lauren Francis, Kirsty Johnstone and Amy Taylor for technical support. Stefan Constantinescu, Mike Simon, Georg Halder kindly provided reagents. Funding was provided by a Senior Cancer Research UK fellowship to MZ and the EU Framework 7 Project ‘Cancer Pathways’.

## References

Arbouzova, N. I. and Zeidler, M. P. (2006). JAK/STAT signalling in Drosophila: insights into conserved regulatory and cellular functions. Development. 133, 2605-2616.

Bach, E. A., Vincent, S., Zeidler, M. P. and Perrimon, N. (2003). A sensitized genetic screen to identify novel regulators and components of the Drosophila janus kinase/signal transducer and activator of transcription pathway. Genetics. 165, 1149-1166.

Bennett, V. and Chen, L. (2001). Ankyrins and cellular targeting of diverse membrane proteins to physiological sites. Curr Opin Cell Biol. 13, 61-67.

Bina, S., Wright, V. M., Fisher, K. H., Milo, M. and Zeidler, M. P. (2010). Transcriptional targets of Drosophila JAK/STAT pathway signalling as effectors of haematopoietic tumour formation. EMBO Rep. 11, 201-207.

Boger, D. L. and Goldberg, J. (2001). Cytokine receptor dimerization and activation: prospects for small molecule agonists. Bioorg Med Chem. 9, 557-562.

Boutros, M., Brás, L. P. and Huber, W. (2006). Analysis of cell-based RNAi screens. Genome Biol. 7, R66.

Brown, R. J. et al. (2005). Model for growth hormone receptor activation based on subunit rotation within a receptor dimer. Nat Struct Mol Biol. 12, 814-821.

Brown, S., Hu, N. and Hombría, J. C. (2003). Novel level of signalling control in the JAK/STAT pathway revealed by in situ visualisation of protein-protein interaction during Drosophila development. Development. 130, 3077-3084.

Cendrowski, J., Mamińska, A. and Miaczynska, M. (2016). Endocytic regulation of cytokine receptor signaling. Cytokine Growth Factor Rev. 32, 63-73.

Chmiest, D. et al. (2016). Spatiotemporal control of interferon-induced JAK/STAT signalling and gene transcription by the retromer complex. Nat Commun. 7, 13476.

Fisher, K. H., Stec, W., Brown, S. and Zeidler, M. P. (2016). Mechanisms of JAK/STAT pathway negative regulation by the short coreceptor Eye Transformer/Latran. Mol Biol Cell. 27, 434-441.

Fisher, K. H., Wright, V. M., Taylor, A., Zeidler, M. P. and Brown, S. (2012). Advances in genome-wide RNAi cellular screens: a case study using the Drosophila JAK/STAT pathway. BMC Genomics. 13, 506.

Gesbert, F., Sauvonnet, N. and Dautry-Varsat, A. (2004). Clathrin-lndependent endocytosis and signalling of interleukin 2 receptors IL-2R endocytosis and signalling. Curr Top Microbiol Immunol. 286, 119-148.

Harrison, D. A., Binari, R., Nahreini, T. S., Gilman, M. and Perrimon, N. (1995). Activation of a Drosophila Janus kinase (JAK) causes hematopoietic neoplasia and developmental defects. EMBO J. 14, 2857-2865.

Heinrich, P. C., Behrmann, I., Haan, S., Hermanns, H. M., Müller-Newen, G. and Schaper, F. (2003). Principles of interleukin (IL)-6-type cytokine signalling and its regulation. Biochem J. 374, 1-20.

Hughes, K. and Watson, C. J. (2012). The spectrum of STAT functions in mammary gland development. JAKSTAT. 1, 151-158.

Kallio, J., Myllymäki, H., Grönholm, J., Armstrong, M., Vanha-aho, L. M., Mäkinen, L., Silvennoinen, O., Valanne, S. and Rämet, M. (2010). Eye transformer is a negative regulator of Drosophila JAK/STAT signaling. FASEB J. 24, 4467-4479.

Kurgonaite, K., Gandhi, H., Kurth, T., Pautot, S., Schwille, P., Weidemann, T. and Bökel, C. (2015). Essential role of endocytosis for interleukin-4-receptor-mediated JAK/STAT signalling. J Cell Sci. 128, 3781-3795.

Liu, J. and Shapiro, J. I. (2003). Endocytosis and signal transduction: basic science update. Biol Res Nurs. 5, 117-128.

Luo, H., Rose, P., Barber, D., Hanratty, W. P., Lee, S., Roberts, T. M., D’Andrea, A. D. and Dearolf, C. R. (1997). Mutation in the Jak kinase JH2 domain hyperactivates Drosophila and mammalian Jak-Stat pathways. Mol Cell Biol. 17, 1562-1571.

Makki, R. et al. (2010). A short receptor downregulates JAK/STAT signalling to control the Drosophila cellular immune response. PLoS Biol. 8, e1000441.

Menning, M. and Kufer, T. A. (2013). A role for the Ankyrin repeat containing protein Ankrd17 in Nod1‐ and Nod2-mediated inflammatory responses. FEBS Lett. 587, 2137-2142.

Mi, H., Huang, X., Muruganujan, A., Tang, H., Mills, C., Kang, D. and Thomas, P. D. (2017). PANTHER version 11: expanded annotation data from Gene Ontology and Reactome pathways, and data analysis tool enhancements. Nucleic Acids Res. 45, D183–;D189.

Michaely, P., Kamal, A., Anderson, R. G. and Bennett, V. (1999). A requirement for ankyrin binding to clathrin during coated pit budding. J Biol Chem. 274, 35908-35913.

Mukherjee, T., Schäfer, U. and Zeidler, M. P. (2006). Identification of Drosophila genes modulating Janus kinase/signal transducer and activator of transcription signal transduction. Genetics. 172, 1683-1697.

Müller, P., Kuttenkeuler, D., Gesellchen, V., Zeidler, M. P. and Boutros, M. (2005). Identification of JAK/STAT signalling components by genome-wide RNA interference. Nature. 436, 871-875.

Murray, P. J. (2007). The JAK-STAT signaling pathway: input and output integration. J Immunol. 178, 2623-2629.

Pasquier, F., Cabagnols, X., Secardin, L., Plo, I. and Vainchenker, W. (2014). Myeloproliferative neoplasms: JAK2 signaling pathway as a central target for therapy. Clin Lymphoma Myeloma Leuk. 14 Suppl, S23-35.

Poulin, F., Brueschke, A. and Sonenberg, N. (2003). Gene fusion and overlapping reading frames in the mammalian genes for 4E-BP3 and MASK. J Biol Chem. 278, 52290-52297.

Qian, H., Buza-Vidas, N., Hyland, C. D., Jensen, C. T., Antonchuk, J., Månsson, R., Thoren, L. A., Ekblom, M., Alexander, W. S. and Jacobsen, S. E. (2007). Critical role of thrombopoietin in maintaining adult quiescent hematopoietic stem cells. Cell Stem Cell. 1, 671-684.

Remy, I., Wilson, I. A. and Michnick, S. W. (1999). Erythropoietin receptor activation by a ligand-induced conformation change. Science. 283, 990-993.

Rossi, F., Charlton, C. A. and Blau, H. M. (1997). Monitoring protein-protein interactions in intact eukaryotic cells by beta-galactosidase complementation. Proc Natl Acad Sci U S A. 94, 8405-8410.

Sansores-Garcia, L., Atkins, M., Moya, I. M., Shahmoradgoli, M., Tao, C., Mills, G. B. and Halder, G. (2013). Mask is required for the activity of the Hippo pathway effector Yki/YAP. Curr Biol. 23, 229-235.

Seubert, N., Royer, Y., Staerk, J., Kubatzky, K. F., Moucadel, V., Krishnakumar, S., Smith, S. O. and Constantinescu, S. N. (2003). Active and inactive orientations of the transmembrane and cytosolic domains of the erythropoietin receptor dimer. Mol Cell. 12, 1239-1250.

Sidor, C. M., Brain, R. and Thompson, B. J. (2013). Mask proteins are cofactors of Yorkie/YAP in the Hippo pathway. Curr Biol. 23, 223-228.

Smith, R. K., Carroll, P. M., Allard, J. D. and Simon, M. A. (2002). MASK, a large ankyrin repeat and KH domain-containing protein involved in Drosophila receptor tyrosine kinase signaling. Development. 129, 71-82.

Staerk, J. and Constantinescu, S. N. (2012). The JAK-STAT pathway and hematopoietic stem cells from the JAK2 V617F perspective. JAKSTAT. 1, 184-190.

Stec, W., Vidal, O. and Zeidler, M. P. (2013). Drosophila SOCS36E negatively regulates JAK/STAT pathway signaling via two separable mechanisms. Mol Biol Cell. 24, 3000-3009.

Tanaka, T., Narazaki, M. and Kishimoto, T. (2014). IL-6 in inflammation, immunity, and disease. Cold Spring Harb Perspect Biol. 6, a016295.

Thomas, C.et al. (2011). Structural linkage between ligand discrimination and receptor activation by type I interferons. Cell. 146, 621-632.

Vainchenker, W. and Constantinescu, S. N. (2013). JAK/STAT signaling in hematological malignancies. Oncogene. 32, 2601-2613.

Vidal, O. M., Stec, W., Bausek, N., Smythe, E. and Zeidler, M. P. (2010). Negative regulation of Drosophila JAK-STAT signalling by endocytic trafficking. J Cell Sci. 123, 3457-3466.

Wang, Y., Tong, X., Li, G., Li, J., Deng, M. and Ye, X. (2012). Ankrd17 positively regulates RIG-I-like receptor (RLR)-mediated immune signaling. Eur J Immunol. 42, 1304-1315.

Zeidler, M. P. and Bausek, N. (2013). The Drosophila JAK-STAT pathway. JAKSTAT. 2, e25353.

